# Performance of convolutional neural networks for identification of bacteria in 3D microscopy datasets

**DOI:** 10.1101/273318

**Authors:** Edouard A Hay, Raghuveer Parthasarathy

## Abstract

Three-dimensional microscopy is increasingly prevalent in biology due to the development of techniques such as multiphoton, spinning disk confocal, and light sheet fluorescence microscopies. These methods enable unprecedented studies of life at the microscale, but bring with them larger and more complex datasets. New image processing techniques are therefore called for to analyze the resulting images in an accurate and efficient manner. Convolutional neural networks are becoming the standard for classification of objects within images due to their accuracy and generalizability compared to traditional techniques. Their application to data derived from 3D imaging, however, is relatively new and has mostly been in areas of magnetic resonance imaging and computer tomography. It remains unclear, for images of discrete cells in variable backgrounds as are commonly encountered in fluorescence microscopy, whether convolutional neural networks provide sufficient performance to warrant their adoption, especially given the challenges of human comprehension of their classification criteria and their requirements of large training datasets. We therefore applied a 3D convolutional neural network to distinguish bacteria and non-bacterial objects in 3D light sheet fluorescence microscopy images of larval zebrafish intestines. We find that the neural network is as accurate as human experts, outperforms random forest and support vector machine classifiers, and generalizes well to a different bacterial species through the use of transfer learning. We also discuss network design considerations, and describe the dependence of accuracy on dataset size and data augmentation. We provide source code, labeled data, and descriptions of our analysis pipeline to facilitate adoption of convolutional neural network analysis for three-dimensional microscopy data.

**Author summary:** The abundance of complex, three dimensional image datasets in biology calls for new image processing techniques that are both accurate and fast. Deep learning techniques, in particular convolutional neural networks, have achieved unprecedented accuracies and speeds across a large variety of image classification tasks. However, it is unclear whether or not their use is warranted in noisy, heterogeneous 3D microscopy datasets, especially considering their requirements of large, labeled datasets and their lack of comprehensible features. To asses this, we provide a case study, applying convolutional neural networks as well as feature-based methods to light sheet fluorescence microscopy datasets of bacteria in the intestines of larval zebrafish. We find that the neural network is as accurate as human experts, outperforms the feature-based methods, and generalizes well to a different bacterial species through the use of transfer learning.

## Introduction

The continued development and widespread adoption of three-dimensional microscopy methods enables insightful observations into the structure and time-evolution of living systems. Techniques such as confocal microscopy [1, 2], two-photon excitation microscopy [3–6], and light sheet fluorescence microscopy [6–12] have provided insights into neural activity, embryonic morphogenesis, plant root growth, gut bacterial competition, and more. Extracting quantitative information from biological image data often calls for identification of objects such as cells, organs, or organelles in an array of pixels, a task that can especially challenging for three-dimensional datasets from live imaging due to their large size and potentially complex backgrounds. Aberrations and scattering in deep tissue can, for example, introduce noise and distortions, and live animals often contain autofluorescent biomaterials that complicate the discrimination of labeled features of interest. Moreover, traditional image processing techniques tend to require considerable manual curation, as well as user input regarding which features, such as cell size, homogeneity, or aspect ratio, should guide and parameterize analysis algorithms. These features may be difficult to know a priori, and need not be the features that lead to the greatest classification accuracy. As data grow in both size and complexity, and as imaging methods are applied to an ever-greater variety of systems, standard approaches become increasingly unwieldy, motivating work on better computational methods.

Machine learning methods, in particular convolutional neural networks (ConvNets), are increasingly used in many fields and have achieved unprecedented accuracies in image classification tasks [13–16]. The objective of machine learning is to use a labeled dataset to train a computer algorithm to make classifications or predictions given new, unlabeled data. Traditional feature-based machine learning algorithms, such as support vector machines and random forests, make use of manually determined characteristics, which in the context of image data could be the eccentricity of objects, their size, their median pixel intensity, etc. The first stages in the implementation of these algorithms, therefore, are the identification of objects by image segmentation methods and the calculation of the desired feature values. In contrast, convolutional neural networks use the raw pixel values as inputs, eliminating the need for determination of object features by the user. Convolutional neural networks use layers consisting of multiple kernels, numerical arrays acting as filters, which are convolved across the input taking advantage of locally correlated information. These kernels are updated as the algorithm is fed labeled data, converging by numerical optimization methods on the weights that best match the training data. ConvNets can contain hundreds of kernels over tens or hundreds of layers which leads to hundreds of thousands of parameters to be learned, requiring considerable computation and, importantly, large labeled datasets to constrain the parameters. Over the past decade, the use of ConvNets has been enabled by advances in GPU technology, the availability of large labeled datasets in many fields, and user-friendly deep learning software such as TensorFlow [17], Theano [18], Keras [19], and Torch [20]. In addition to high accuracy, ConvNets tend to have fast classification speeds compared to traditional image processing methods. There are drawbacks, however, to neural network approaches. As noted, they require large amounts of manually labeled data for training the network. Furthermore, their selection criteria, in other words the meanings of the kernels’ parameters, are not easily understandable by humans [21].

There have been several notable examples of machine learning methods applied to biological optical microscopy data [22, 23], including bacterial identification from 2D images using deep learning [24],pixel-level image segmentation using deep learning [25–27], subcellular protein classification [28], and detection of structures within C. elegans from 2D projections of 3D image stacks using support vector machines [29]. Nonetheless, it is unclear whether ConvNet approaches are successful for thick, three-dimensional microscopy datasets, whether their potentially greater accuracy outweighs the drawbacks noted above, and what design principles should guide the implementation of ConvNets for 3D microscopy data.

To address these issues, we applied a deep convolutional neural network to analyze three-dimensional light sheet fluorescence microscopy datasets of gut bacteria in larval zebrafish (Fig 1 a,b) and compared its performance to that of other methods. These image sets, in addition to representing a major research focus of our lab related to the aim of understanding the structure and dynamics of gut microbial communities [10, 30–32], serve as exemplars of the large, complex data types increasingly enabled by new imaging methods. Each 3D image occupies roughly 5 GB of storage space and consists of approximately 300 slices separated by 1 micron, each slice consisting of 6000 × 2000 pixel 2D images (975×325 microns). These images include discrete bacterial cells, strong and variable autofluorescence from the mucus-rich intestinal interior [33], autofluorescent zebrafish cells, inhomogeneous illumination due to shadowing of the light sheet by pigment cells, and noise of various sorts. The bacteria examined here exist predominantly as discrete, planktonic individuals. Other species in the zebrafish gut exhibit pronounced aggregation; identification of aggregates is outside the scope of this work, though we note that the segmentation of aggregates is much less challenging than identification of discrete bacterial cells, due to their overall brightness and size. The goal of the analysis described here is to correctly classify regions of high intensity as bacteria or as non-bacterial objects.

**Fig 1.**
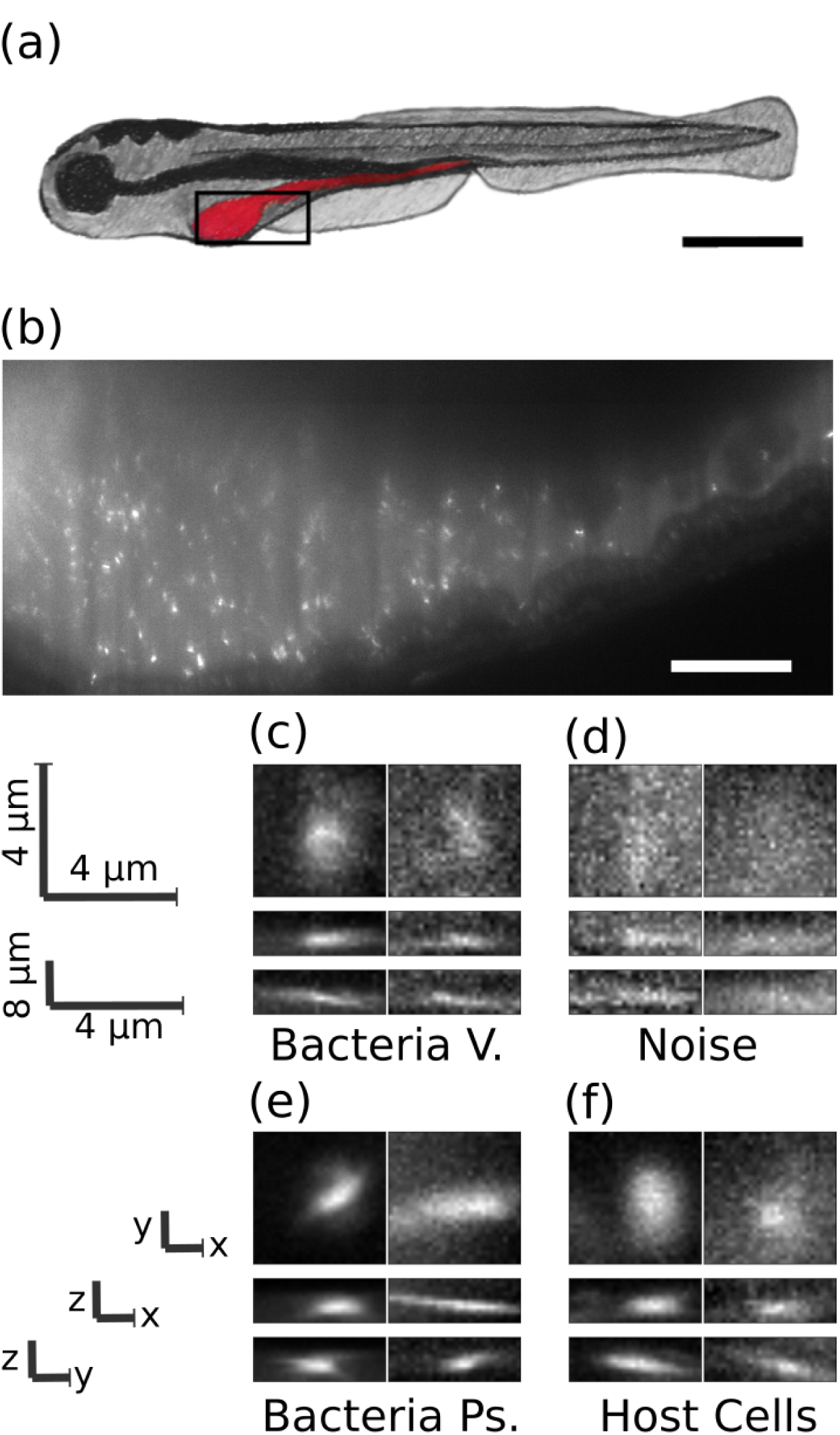
Images of bacteria in the intestine of larval zebrafish. a) Schematic illustration of a larval zebrafish with the intestine highlighted in red. Scale bar: 0.5 mm. b) Single optical section from light sheet fluorescence microscopy of the anterior intestine of a larval zebrafish colonized by GFP expressing bacteria of the commensal Vibrio species ZWU0020. Scale bar: 50 microns. c) z, y and x projections from 28×28×8 pixel regions of representative individual Vibrio bacteria, d) non-bacterial noise, e) individual bacteria of the genus Pseudomonas, species ZWU0006, and f) autofluorescent zebrafish cells.

Using multiple testing image sets, we compared the performance of the convolutional neural network to that of humans as well as random forest and support vector machine classifiers. In brief, the ConvNet’s accuracy is similar to that of humans, and it outperforms the other machine classifiers in both accuracy and speed across all tested datasets. In addition, the ConvNet performs well when applied to planktonic bacteria of a different genus through the use of transfer learning, in which partial transference of network weights dramatically lowers the amount of new labeled data that is required. We explored the impacts on the ConvNet’s performance of network structure, the degree of data augmentation using rotations and reflections of the input data, and the size of the training data set, providing insights that will facilitate the use of ConvNets in other biological imaging contexts.

Analysis code as well as all ∼ 21, 000 manually labeled 3D image regions-of-interest are provided; see Methods for details and urls to data locations.

## Results

### Data

The image data we sought to classify consist of three-dimensional arrays of pixels obtained from light sheet fluorescence microscopy of bacteria in the intestines of larval zebrafish [10, 30–32]. Fig 1B shows a typical optical section from an initially germ-free larval zebrafish, colonized by a single labeled bacterial species made up of discrete, planktonic individuals expressing green fluorescent protein; a three-dimensional scan is provided as Supplementary Movie 1. All the data assessed here were derived from fish that were reared germ free (devoid of any microbes) [34] and then either mono-associated with a commensal bacterial species or left germ free. Nine scans are of fish mono-associated with the commensal species ZWU0020 of the genus Vibrio [10, 35, 36], two scans are of fish in which the zebrafish remained germ-free, and a single scan is from a fish mono-associated with Pseudomonas ZWU0006 [31]. For each 3D scan, we first determined the intestinal space of the zebrafish using simple thresholding and detected bright objects (“blobs”) using a difference of Gaussians method described further in Methods. From each blob, we extracted 28×28×8 pixel arrays (4.5×4.5×8 microns), which served as the input data to the neural network, to be classified as bacterial or non-bacterial.

Since there is no way to obtain ground truth values for bacterial identity in images, we manually classified blobs to serve as the training data for the neural network, using our expertise derived from considerable prior work on three dimensional bacterial imaging. Notably, in prior work we showed that the total bacterial abundance determined by manually corroborated feature-based bacterial identification from light sheet data corresponds well with the total bacterial abundance as measured through gut dissection and serial plating assays [30]. In Fig 1C-F we show representative images of blobs corresponding to bacteria and noise.

In order to estimate an upper bound on the classification accuracy we can expect from the learning algorithms, we chose a single image scan which we judged to be typical of a noisy, complex 3D image of the intestine of a larval zebrafish colonized by bacteria. We then had six lab members with considerable light sheet microscopy experience individually label each of the detected potential objects as either a bacterium or not. We show in Fig 2A the agreement between lab members. Excluding human 3 the agreement between any pair of humans is always above 0.87. The outlier, human 3, is the person with the least experience with the imaging data, namely the principal investigator.

**Fig 2.**
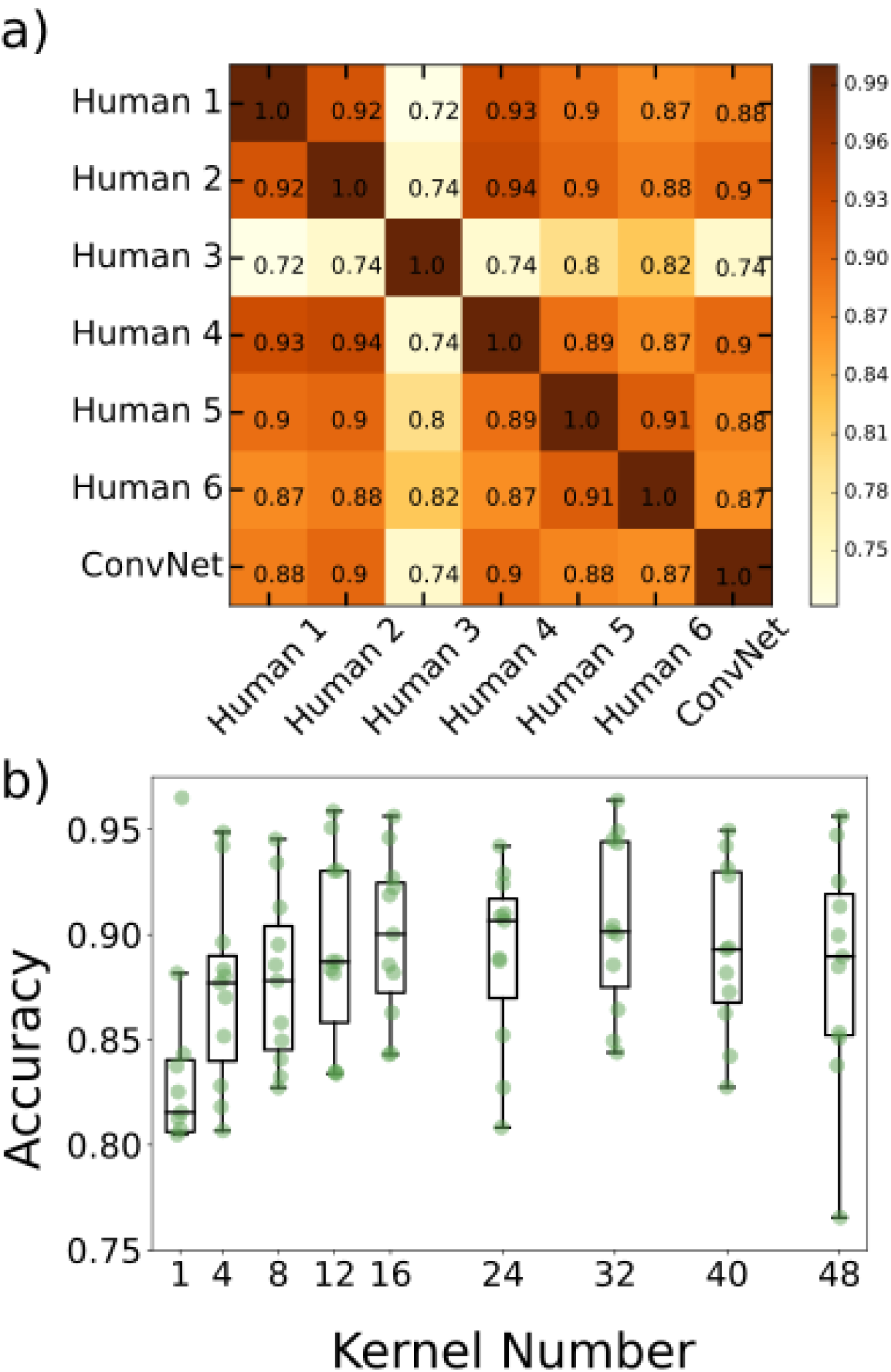
Creation of the 3-D convolutional neural network. a) Agreement matrix between six individuals (members of the authors’ research group), evaluated on a single dataset of images of Vibrio bacteria, and between those humans and the convolutional neural network. b) Accuracy vs number of kernels per layer using cross validation across the various imaging datasets, where the x-axis denotes the number of kernels in the first convolutional layer. The second convolutional layer for each plotted point has twice as many kernels as the first.

We next created a set of labeled data by manual classification of blobs from the 9 Vibrio scans and 2 scans of germ-free fish, consisting in total of over 20,000 objects. Including scans from germ-free fish is particularly important to enable accurate counting of low numbers of bacteria, which arise naturally due to extinction events [10] and population bottlenecks [35].

### Network Architecture

As detailed in Methods, we used Google’s open-source Tensorflow framework [2016, Abadi] to create, test, and implement 3D convolutional neural networks. Such networks have many design parameters and options, including the number, size, and type of layers, the kernel size, the downsizing of convolution output by pooling, and parameter regularization. In general, overly small networks can lack the complexity to characterize image data, though their limited parameter space is less likely to lead to overfitting. Conversely, larger networks can tackle more complex classification schemes, but demand more training data to constrain the large number of parameters, and also carry a greater computational load. In between these extremes, many design variations will typically give similar classification accuracy. We chose a simple architecture consisting of two convolutional layers followed by a fully connected layer. The first and second convolutional layers contain 16 and 32 5×5×2 kernels, respectively. Each layer is followed by 2×2×2 max pooling as further described in Methods. The final layer is a fully connected layer consisting of 1024 neurons with a dropout rate of 0.5 during training. After this, softmax regression is used for binary classification.

We explored various alterations of our network architecture, and illustrate here the effect of simply varying the number of kernels per convolutional layer. We assessed the classification accuracy as a function of the number of kernels in layer 1, with the number of kernels in layer 2 being double this. Accuracy was calculated using cross validation, training on all but one image dataset (where an image dataset is a complete three-dimensional scan of the gut of one zebrafish), testing on the remaining image dataset, and repeating with different train/test combinations. The network accuracy initially increases with kernel number and plateaus at roughly 16 kernels, beyond which the variance in accuracy increases (Fig 2B). Therefore, increasing the number of kernels beyond approximately 16 gives little or no improvement in accuracy at the expense of model complexity and increased variability.

### Network Accuracy Across Image Datasets

We trained the ConvNet using manually labeled data from eight of the Vibrio image datasets and the two datasets from germ-free fish (devoid of gut bacteria) and then tested it on the remaining manually labeled Vibrio image dataset that was used to assess inter-human variability, described above. The agreement between the neural network and humans (mean ± std. dev. 0.89 ± 0.01) was indistinguishable from the inter-human agreement (mean ± std. dev. 0.90 ± 0.02), again excluding human 3, indicating that the ConvNet achieves the practical maximum of bacterial classification accuracy (Fig 2A). Examples of images for which all humans agreed on the classification, and in which there was disagreement, are provided in the Supplementary Text.

To further test the network’s consistency across different imaging conditions we applied it separately to each of the 3D image datasets of larval zebrafish intestines. We also tested, with the same procedure and data, random forest and support vector machine classifiers to address the question of whether or not the ConvNet outperforms typical feature based learning algorithms. We first consider two experiment types: zebrafish intestines mono-associated with Vibrio ZWU0020 (9 image datasets, i.e. 9 complete three-dimensional scans from of different zebrafish) and germ-free zebrafish (2 image datasets). Classifier accuracy for each Vibrio-colonized or empty-gut image scan was determined by cross-validation, training the network using all of the other image datasets, and testing on the dataset of interest. To test the variance in accuracy due to the training process, we performed three repetitions of each train/test combination using the same data. We found that the neural network outperforms the feature based algorithms on every image dataset (Fig 3), and also shows less variation in accuracy between the different datasets. The enhanced accuracy from the neural network is especially dramatic for germ-free datasets, for which it achieves over 90% accuracy, in contrast to less than 75% for feature based methods. For a given test dataset, the training variance for the convolutional neural network is small but nonzero, indicating that the network training algorithm finds similar, but not identical, minima with different (random) initializations on the same training data. It is also small for the random forest classifier. Interestingly, it is zero for the SVM classifier, indicating that given the same dataset, the algorithm is finding the same minimum.

**Fig 3.**
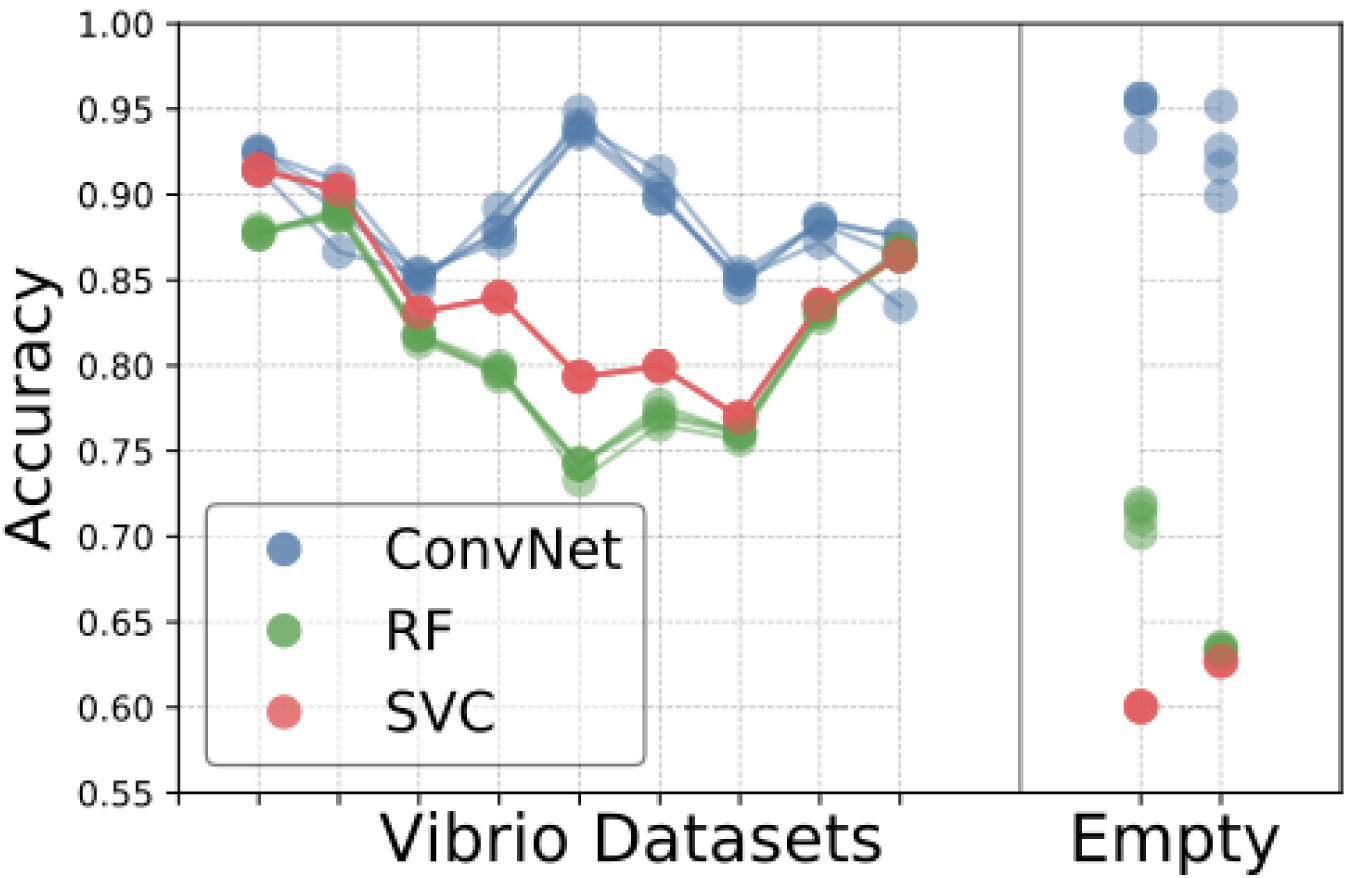
Comparison of Convnet and feature based learning algorithms across all datasets. Comparison of accuracies for the various learning algorithms (convolutional neural network, support vector classifier, and random forest) across different Vibrio image datasets, as well as two image datasets from fish devoid of gut bacteria. Each accuracy was determined by training on the data from all of the other datasets, and testing on the dataset of interest.

The random forest, support vector machine, and neural network classifiers process roughly 300, 400, and 950 images per second, respectively; i.e. the neural network runs 2-3 times faster than the feature based learning algorithms on the same data.

### Training Size and Data Augmentation

Convolutional Neural Networks famously require large amounts of training data which must often, as is the case here, be evaluated and curated by hand. To assess the scale of manual classification required for good algorithm performance, which is a key issue for future adoption of neural networks in biological image analysis, we explored the effect on the network’s accuracy of varying the amount of training data. We set aside 25% of the images from each of the Vibrio and germ-free fish image scans and trained the network using an increasing number of images from the remaining data. We increased the amount of training data in two different ways. First, we consecutively added to the training set all images from each image dataset excluding a subset of the images previously reserved for testing (labeled Test: new datasets in Fig 4A). Second, we randomly shuffled the training images from all the image scans, adding 1500 images to the training set over each iteration (labeled Train/test split in Fig 4A). For the first method, enlargement of the training set corresponds to a greater amount of data as well as data from more diverse biological sources. For the second, data size increases but the biological variation sampled is held constant. In both cases, accuracy plateaus at a number of images on the order of 10,000 (Fig 4A). The rise in accuracy with increasing training data size is only slightly more shallow with the first method, surprisingly, demonstrating that within-sample variation is sufficient to train the network.

**Fig 4.**
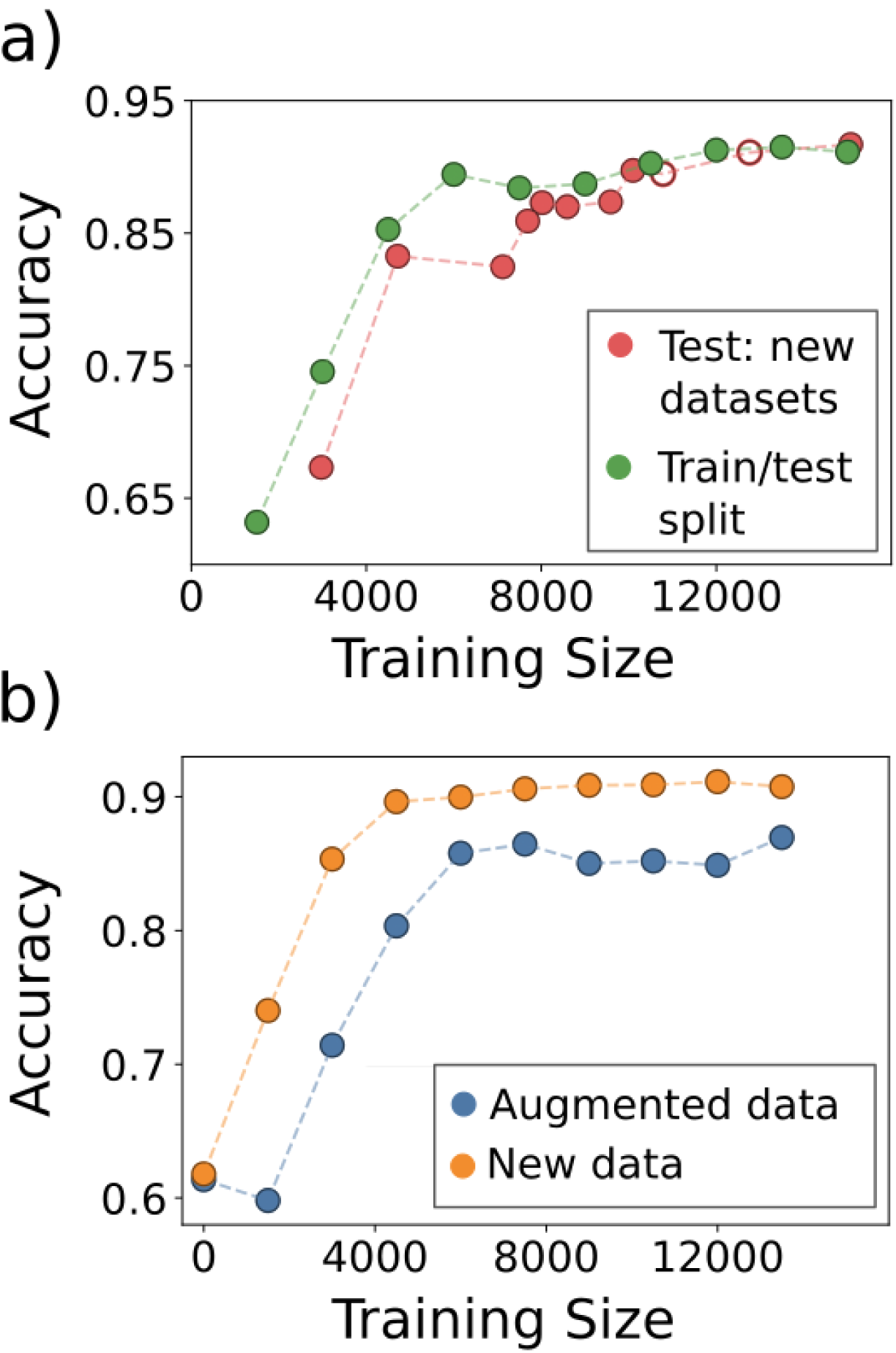
Data augmentation. Examining the accuracy of the CNN as a function of a) varying the training data size by adding images from biologically distinct datasets (Test: new datasets) or by adding images randomly from the full set of images (Train/test split), and b) transformation of the data by image rotations and reflections. In (a), the two empty circles represent the inclusion of the datasets from empty (germ-free) zebrafish intestines.

Data augmentation, the alteration of input images through mirror reflections, rotations, cropping, and the addition of noise, etc., is commonly used in machine learning to enhance training dataset size and enable robust training of neural networks. To characterize the utility of data augmentation for 3D bacterial images, we focused in particular on image rotations and reflections, because the bacteria have no preferred orientation and hence augmentation by these methods creates realistic training images. We note that data augmentation is not necessary for feature based learning methods in which parity and rotational invariance can be built into the features used for classification. Obviously, augmented data is not independent of the actual training data, and so does not supply wholly new information. We were curious as to how including rotated and reflected versions of previously seen data compares, in terms of network performance, to adding entirely new data, a comparison that is useful if evaluating the necessity of performing additional imaging experiments. To test this, we compared the accuracies of the network when adding new data to that when adding rotated and reflected versions of existing data. We started with a fixed number of 1500 total objects randomly sampled from the entire set and, in the case of including new data, added another random 1500 objects at each iteration. For the augmented data, we applied random rotations and reflections to the original 1500 objects to iteratively increase the training size by 1500 objects. Each trained network was tested on the same test set of objects as that of Fig 4A. As shown in Fig 4B, the addition of new data leads to a plateau in accuracy of roughly 90% while for augmented data the plateau value is around 88%. This result demonstrates that, in the context of our network, simply augmenting existing data can raise classification accuracy to nearly the optimal level achieved by new, independent data.

### Transfer Learning

We assessed the accuracy of the convolutional neural network on images of discrete gut bacteria of another species, of the genus Pseudomonas. Training solely on the Vibrio images and testing on Pseudomonas gives ∼ 75% accuracy (Fig 5). However, this is much lower than the ∼ 85 − 95% accuracy obtained on Vibrio images (Fig 4); the Pseudomonas species is not an exact morphological mimic of the Vibrio species. The Pseudomonas dataset is small (1190 images); using 80% of its images for de novo neural network training gives ∼ 72% accuracy in identifying Pseudomonas in test datasets (Fig 5). We suspected that the general similarity of each species as rod-like, few-micron-long cells would allow transfer learning, in which a model trained for one task is used as the starting point for training for another task [37, 38]. Using the network weights from training on Vibrio image datasets, as before, as the starting values for training on the small Pseudomonas dataset gives over 85% accuracy in classifying Pseudomonas (Fig 5).

**Fig 5.**
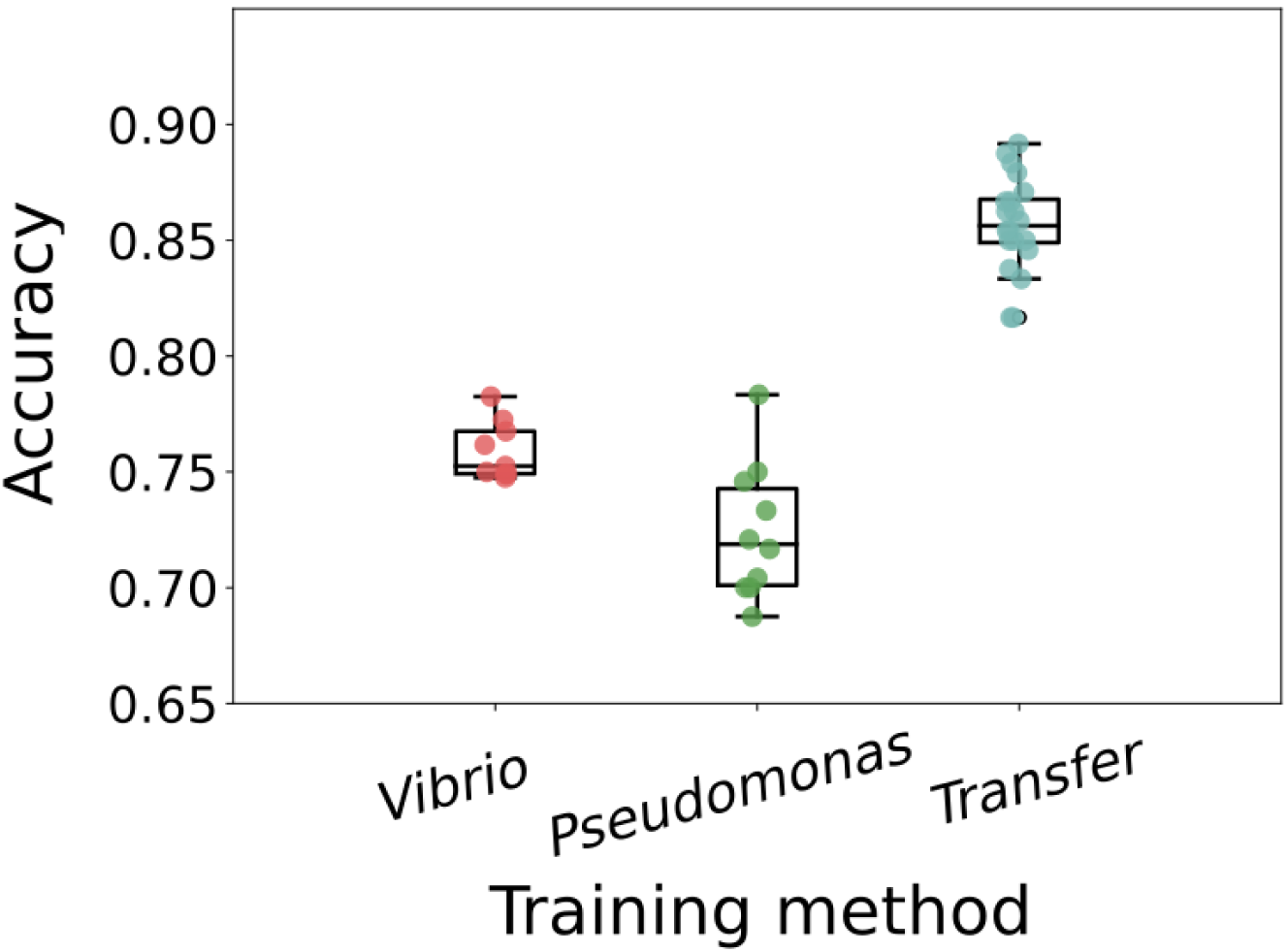
Transfer Learning on New Bacterial Species. The accuracy of Pseudomonas classification with convolutional neural networks trained in different ways. “Vibrio” indicates training on images of Vibrio bacteria, “Pseudomonas” indicates training on the small Pseudomonas image dataset, and “Transfer” indicates using the Vibrio-derived network weights as the starting point for training on Pseudomonas images. For training only on Vibrio images, the different data points come from random weight initialization, random data ordering, and random augmentation. For training only on Pseudomonas images, and for transfer learning, the different data points are from random train/test splits of the Pseudomonas data.

## Discussion

We find that a 3D convolutional neural network for binary classification of bacteria and non-bacterial objects in 3D microscopy data of the larval zebrafish gut yields high accuracy without unreasonably large demands on the amount of manually curated training data. Specifically, the convolutional neural network obtains human-expert-level accuracy, runs 2-3 times faster than other standard machine learning methods, and is consistent across different datasets and across planktonic bacteria from two different genera through the use of transfer learning. It reaches these performance metrics after training on fewer than 10,000 human-classified images, which require approximately 20 person-hours of manual curation to generate. Moreover, augmented data in the form of rotations and reflections of real data contributes effectively to network training, further reducing the required manual labor. Experiments of the sort presented here typically involve many weeks of laboratory work. Neural network training, therefore, is a relatively small fraction of the total required time.

In many biological imaging experiments, including our own, variety and similarity are both present. Multiple distinct species or cell types may exist, each different, but with some morphological similarities. It is therefore useful to ask whether such similarities can be exploited to constrain the demands of neural network training. The concept of transfer learning addresses this issue, and we find that applying it to our bacterial images achieves high accuracy despite small labeled datasets, an observation that we suspect will apply to many image-based studies.

Though the data presented here came from a particular experimental system, consisting of fluorescently labeled bacterial species within a larval zebrafish intestine imaged with light sheet fluorescence microscopy, they exemplify general features of many contemporary three-dimensional live imaging applications, including large data size, high and variable backgrounds, optical aberrations, and morphological heterogeneity. As such, we suggest that the lessons and analysis tools provided here should be widely applicable to microbial communities [39] as well as eukaryotic multicellular organisms.

We expect the use of convolutional neural networks in biological image analysis to become increasingly widespread due to the combination of efficacy, as illustrated here, and the existence of user-friendly tools, such as TensorFlow, that make their implementation straightforward. We can imagine several extensions of the work we have described. Considering gut bacteria in particular, extending neural network methods to handle bacterial aggregates is called for by observations of a continuum of planktonic and aggregated morphologies [31]. Considering 3D images more generally, we note that the approach illustrated has as its first step detection of candidate objects (“blobs”), which requires choices of thresholding and filtering parameters. Alternatively, pixel-by-pixel segmentation is in principle possible using recently developed network architectures [13, 40], which could enable completely automated processing of 3D fluorescence images. In addition, pixel-based identification of overall morphology (for example, the location of the zebrafish gut) could further enhance classification accuracy, by incorporating anatomical information that constrains the possible locations of particular cell types.

## Methods

### Light Sheet Microscopy Image Data

Three-dimensional scans of the intestines of larval zebrafish, derived germ-free and colonized by fluorescently labeled bacteria prior to imaging, were obtained using light sheet fluorescence microscopy as described in Refs. [10, 30, 31]. All experiments involving zebrafish were carried out in accordance with protocols approved by the University of Oregon Institutional Animal Care and Use Committee.

The microscope was based on the design from Keller et al [6], and has been described elsewhere [30, 39]. In brief: a laser is rapidly oscillated creating a thin sheet of light used to illuminate a section of the specimen, in this case, a larval zebrafish. An objective lens is seated perpendicular to the laser sheet, focusing two-dimensional images onto a sCMOS camera. The specimen is scanned through the sheet along the detection axis, thereby constructing a 3D image. The camera exposure time was 30 ms, and the laser power of the laser was 5 mW as measured between the theta-lens and excitation objective.

Of the twelve image datasets used for this work, nine were of the zebrafish commensal bacterium Vibrio sp. ZWU0020, one was of a Pseudomonas commensal sp. ZWU0006, and two were from germ-free fish, devoid of any bacteria.

An example 3D image dataset of the anterior “bulb” of one larval zebrafish gut is available at the link noted in the README.md file at github: https://github.com/rplab/Bacterial-Identification, together with the 6 lab members’ labels for each detected object in the volume, the convolutional neural network’s classification, and each of the extracted region-of-interest voxels. Other image sets are available upon request; for each zebrafish gut, the full image dataset is roughly 1 GB in size.

### Segmentation and Blob Detection

Rough segmentation of the intestine was performed using histogram equalization of each individual z-stack followed by a moving average over 30 consecutive images in the z-stack followed by hard thresholding to create a binary mask that overestimated the size of the intestine. While extremely rough, this technique requires no manual editing or outlining. After this, blob detection was performed using the difference of Gaussians technique from the scikit-image library on each two-dimensional image, and the blobs were linked together across consecutive images in each stack. Regions 28×28×8 pixels in size centered at each detected blob were then saved to be labeled by hand as either a bacterium or noise. The code for extracting the regions of interest is publicly available on Github at https://github.com/rplab/Bacterial-Identification.

From the 12 datasets, 20,929 images were hand labeled of which 38% were bacteria and 62% were noise. Hand labeling took roughly 1-2 hours per scan. All of the 28×28×8 pixel images and the corresponding labels are available from links in the README.md file at the Github repository https://github.com/rplab/Bacterial-Identification.

All code for the project was written in Python.

### Random Forest and Support Vector Machine Classifiers

Over sixty features were created initially. These were assessed using scikit-learn’s feature_importances_, from which the thirty one most helpful features were retained. The features used included geometric properties obtained by ellipse-fitting and texture-based characteristics; a detailed list is provided in the python code features.pyprovided on Github: https://github.com/rplab/Bacterial-Identification. The data were tested using both a random forest and support vector classifier from the scikit-learn library. The random forest used 500 estimators. The support vector classifier from sci-kit learn, sklearn.svm.SVC(), was tested over a range of parameters and kernels using scikit-learn’s GridSearchCV which yielded highest accuracy when using a radial basis function kernel with penalty C=1.

### Convolutional Neural Network

The 3D convolutional neural network was created using Google’s TensorFlow. Each input image was 28×28×8 pixels. The network consisted of two convolutional layers followed by a fully connected layer. The first layer was composed of 16, 5×5×2 kernels of stride 2 and same padding followed by 2×2×2 max pooling, the second layer contained 32 5×5×2 kernels of the same stride and padding and was also followed by 2×2×2 max pooling. We chose to double the number of kernels after max pooling as in [41]. After the final convolutional layer we employed a fully connected layer consisting of 1024 neurons. The classes were then determined using a softmax layer. The network had a dropout of 0.5, a learning rate of 0.0001 and the data was trained over 120 epochs randomly rotating and reflecting each image over each epoch unless otherwise specified. The weights were updated using the Adam optimization method and we use leaky-ReLu activation functions. During each epoch of training, each input image has a fifty percent probability of receiving a reflection in x, y and z followed by a fifty percent probability of subsequently being transposed. This particular scheme was chosen due to its low computational load. We have made the code for this convolutional neural network is available on Github at https://github.com/rplab/Bacterial-Identification.

### Computer Specs and Timing

The code was implemented on using python 3.5 on Ubuntu 16.04, with a Intel Core i7-4790 CPU with an Nvidia GeForce GTX 1060 graphics card on a computer with 32 GB of RAM. With this hardware it took roughly one minute to train and create the features for the RF and SVC using about 17,000 images, and roughly one hour to train the 3D ConvNet on the same number of images.

## Acknowledgments

We thank Rose Sockol and the University of Oregon Zebrafish Facility staff for fish husbandry, Sophie Sichel for preparation of germ-free zebrafish, and many members of the authors’ research group for useful comments and conversations.

